# Rapid appetitive transitions are sculpted by amygdala to accumbens pathways

**DOI:** 10.1101/2021.04.11.435530

**Authors:** E. Zayra Millan, Jun Hua Lim, John M. Power, Gavan P. McNally

**Author notes:** Correspondence to: E. Zayra Millan, PhD, School of Psychology, UNSW Sydney NSW 2052, Sydney Australia, E.-mail. **Author contributions** <u>Conceptualization</u>: EZM, JMP, GPM. <u>Experimentation</u>: EZM, JL, JMP. <u>Formal Analysis</u>: EZM, JMP. <u>Writing – original draft:</u> EZM, JMP, GPM. <u>Writing – Review & Editing</u>: All authors. <u>Funding acquisition</u>: GPM, JMP.

## Abstract

Foraging, pursuit, and predation rapidly transition into behavioral quiescence during reward capture and consumption. While appetitive-consummatory dissociations are embedded at both psychological and neural levels, the mechanisms controlling switches or transitions between appetitive seeking and consummatory behaviors remain poorly understood. Here we identify the BLA→AcbSh pathway as critical to these transitions by showing that this pathway inhibits the appetitive seeking response in the presence of consummatory demands. Using an appetitive cue-discrimination task in male rats, we show that reward delivery is a significant driver of seeking inhibition and that a BLA→AcbSh pathway mediates this inhibition. This role in suppressing seeking responses during periods of consumption was not due to a general suppression of behavior because responding to other cues during the same test was unaffected. Moreover, it was specific to the BLA→AcbSh pathway, because the contribution of the BLA→AcbC pathway to appetitive switching was distinct and modest. State-dependent silencing of BLA→AcbSh revealed that the modulation of seeking before and after reward delivery are co-dependent. Finally, we found that BLA terminals in AcbSh have functional connectivity to LH-projecting AcbSh neurons, thereby identifying a BLA→AcbSh→LH pathway as a putative route for the rapid regulation of appetitive behaviors. Taken together, these findings suggest that the BLA→AcbSh pathway is a core component of an appetitive switching system, recruited under conditions requiring rapid or dynamic shifts in appetitive behavior, and that this pathway enables these shifts by actively inhibiting seeking.

**Significance statement:** Foraging, pursuit, and predation quickly transition into behavioral quiescence during reward capture and consumption. These transitions are critical for flexible and responsive sequences of behavior. Here we show that behavioral transitions are actively controlled at the limbic-striatal interface. We identify reward receipt as a proximal trigger for transition between reward seeking and taking, we identify active inhibition as the functional operation of this transition, and we identify the basolateral amygdala→accumbens shell pathway as critical to this functional operation.

---

Transitions between different behavioral repertoires, such as between seeking and consuming reward, are essential to dynamic, adaptive behavior. How do these behavioral transitions occur? Mechanical transitions, such as those used in bicycles, require the selection and engagement of active gear states. In contrast, electronic circuits require termination of one state as a necessary step for switching to another (e.g., S-R latches and flip flops). The neurobiological processes that mediate transitions in motivated behavior are poorly understood and it remains unclear whether they require selection of simultaneously active states or the suppression of one for the emergence of the other.

Previous studies describing the dichotomy between seeking a reward and consuming it have long recognized that distinct neurobiological mechanisms mediate appetitive (i.e. approach) versus consummatory behaviors (Ikemoto & Panksepp, 1999; Pfaus & Phillips, 1991). Recent findings show that this separation is particularly notable in lateral hypothalamus (LH) where distinct neuronal populations encode appetitive versus consummatory behavior (Jennings, Rizzi, Stamatakis, Ung, & Stuber, 2013). Yet, how these distinct appetitive and consummatory behavioral modes are linked–a basic requirement for organized sequences of adaptive behavior–remain poorly understood.

The transition from reward seeking (via approach) to taking (via consumption) requires significant behavioral change. It is predicated upon close coupling of appetitive and consummatory behavioral sequences. Specifically, this transition requires initiation of consummatory or taking behaviors defined by the precise properties of the reward (e.g., ingesting, handling, hoarding) and the concurrent cessation of any seeking behaviors (e.g., approach, foraging, predation, pursuit) that would otherwise interfere with this consumption, and finally it requires termination of motivational drive (e.g. hunger) underpinning behavior (Allen et al., 2019; Hilliard, Domjan, Nguyen, & Cusato, 1998; Ikemoto & Panksepp, 1999; Konorski, 1967; Timberlake, 1994).

From this perspective, the nucleus accumbens (Acb), and its major glutamatergic input from the basolateral amygdala (BLA), may play a crucial role in these transitions. The Acb is implicated in suppression of appetitive seeking behaviors in humans (Kruse, Tapia Leon, Stark, & Klucken, 2017) while in other animals, increases in basal Acb activity are associated with inhibition of seeking responses (German & Fields, 2007; Ghazizadeh, Ambroggi, Odean, & Fields, 2012). Moreover, facilitating excitatory transmission in Acb medial shell (AcbSh) enhances inhibition of reward seeking (Sutton et al., 2003) whereas facilitating inhibition in AcbSh disinhibits reward seeking (Ambroggi, Ghazizadeh, Nicola, & Fields, 2011; Di Ciano, Robbins, & Everitt, 2008; Floresco, McLaughlin, & Haluk, 2008; Lafferty, Yang, Mendoza, & Britt, 2020; Millan, Furlong, & McNally, 2010; Millan & McNally, 2011; Millan, Reese, Grossman, Chaudhri, & Janak, 2015; Peters, LaLumiere, & Kalivas, 2008). This role is strongly associated with AcbSh (Ambroggi et al., 2011; Peters et al., 2008) and its GABAergic innervation of LH (Gibson et al., 2018; O’Connor et al., 2015). The BLA is a major source of excitatory transmission in Acb (Britt et al., 2012; MacAskill, Cassel, & Carter, 2014; Yu et al., 2017). Although this projection mediates the facilitatory impact of reward cues on seeking (Ambroggi et al., 2011; Beyeler et al., 2016; Everitt et al., 1999; Namburi et al., 2015) it also plays a role in inhibiting seeking because activating this pathway terminates reward seeking (Millan, Kim, & Janak, 2017) and silencing or disconnecting this pathway can disinhibit reward seeking (Lafferty et al., 2020; Millan & McNally, 2011).

Here we investigated the mechanisms underlying rapid transitions from appetitive seeking to consumption. We show that reward delivery is a key inflection point in this behavioral sequence, that reward delivery actively suppresses further seeking behavior, that this suppression is mediated via a BLA→AcbSh pathway, and that this pathway controls LH-projecting AcbSh neurons. Together, our findings show that the BLA→AcbSh pathway is fundamental to the rapid regulation of appetitive behaviors and that this role may be essential for linking appetitive seeking with consummatory taking.

## Materials and Methods

### Subjects

Experimentally naive, male Sprague Dawley (*in vivo* optogenetics, *Experiments 3 and 4*) (ARC, Perth, Western Australia) or Long-Evans rats (remaining experiments, UNSW, Randwick; 6-8 weeks at arrival) were group housed (4 per cage) in ventilated polycarbonate cages in a temperature (21°C) and light-regulated vivarium (lights on 7am, 12 h light/dark cycle) with partial enrichment provided by key rings, wood blocks or tunnels. Water was freely available throughout the duration of these studies. Rats were maintained at 90% of their bodyweight for *in vivo* studies during training and testing. Experiments were approved by the UNSW Animal Care and Ethics Committee and performed in accordance with the Animal Research Act 1985 (NSW), under the guidelines of the National Health and Medical Research Council Code for the Care and Use of Animals for Scientific Purposes in Australia (2013).

### Apparatus

Behavioral training and testing were conducted in Med-Associates (St. Albans, VT, USA) chambers [24 cm (length) x 30 cm (width) x 21 cm height)] enclosed in ventilated, sound-attenuating cabinets [59.5 cm (length) x 59 cm (width) x 48cm (height)]. A single recessed magazine receptacle was fitted in the center panel of a sidewall, attached to a pellet delivery system that delivered 45g grain pellets (Bio-Serv, Flemington, N.J. USA) into the dish. 2 x grain pellets served as the unconditioned stimulus (US). A speaker attached to the left-side wall of the chamber was used to deliver auditory conditioning stimuli (CSs). An incandescent stimulus light (white) was positioned on either side of the magazine to deliver the visual CS.

### AAV Constructs and retrograde tracing

We used AAV5-serotyped AAV vectors. Halorhodospin (eNpHR3.0; AAV5/CaMKII-eNpHR3.0-eYFP; ∼10^12^ infectious units ml^−1^); channelrhodopsin (ChR2, AAV5/CaMKII-ChR2-eYFP; ∼10^12^ infectious units ml^−1^) and yellow fluorescent protein (eYFP; AAV5/CaMKII-eYFP or AAV5-DIO-eYFP) were packaged by the University of North Carolina Vector Core (Chapel Hill, NC, USA). AAVs were placed under the calcium-calmodulin kinase IIα (CaMKIIα) promoter to target pyramidal projections in the BLA. For retrograde tracing, animals were injected with fluorescent cholera toxin subunit B (CTb-AlexaFluor488, CTb-AlexaFluor594; ThermoFisher, C34775 and C34777, respectively) into AcbSh and AcbC (counterbalanced).

### Surgery

Rats were anaesthetized via 1.3ml/kg ketamine (Ketamil, Ilium) (100 mg/ml) and 0.3 mg/kg xylazine (Xylazil, Ilium) (20 mg/ml.) They received s.c carprofen (Rimadyl, Zoetis) and 0.5% bupivacaine under the incision site. For the *in vivo* optogenetics experiments we infused BLA bilaterally with AAVs encoding eNpHR3.0 or the control transgene for eYFP. BLA coordinates from bregma were: AP: −2.85; ML: ±4.85; DV: −8.85. AAV injections were delivered in a volume of 0.5µl, at a rate of 0.1ul min^−1^. Injections were left in place for an additional 10 min to allow diffusion of viral particles away from the injection site. Fiber optic cannulae (400µm core diameter) attached to 12.7mm (2.5mm ⌀) stainless steel or ceramic ferrules were bilaterally implanted above AcbSh or core AcbC using coordinates AP: +1.35; ML for AcbSh: ±1.7 (6° angle); ML for AcbC: ±2.7 (6° angle); DV: −6.55 (AcbSh) or −6.60 (AcbC). For the anatomical tract-tracing study, rats received ipsilateral injections of 0.2µl CTb-AlexaFluor488 and CTb-AlexaFluor594, one in each of AcbSh and AcbC (counterbalanced) at a rate of .1ul min^−1^.

For *in vitro* electrophysiology, we combined optogenetic stimulation with retrograde tract-tracing. Retrograde tracer Ctb-AlexaFluor555 was injected unilaterally into the LH and AAV containing ChR2 injected into the BLA using coordinates described above.

Rats were permitted to recover from surgery in their homecages for a minimum of 14 days prior to any experimentation. During this time, rats had free access to food and water and were monitored by the experimenters.

### Procedure

#### Pavlovian appetitive conditioning

Rats received a single 64 min magazine-training session comprising 16 x food pellet deliveries into the magazine at a variable time (VT) of 240s. Subsequently, rats received daily Pavlovian conditioning sessions (8d for behavioral studies; 10d for optogenetic studies) involving presentations of three types of cues. CS1 and CS2 served as auditory conditioning stimuli (CS+, 1800 Hz tone or clicker, 10s duration, 85dB, counterbalanced). Each was paired with two pellets, delivered 2s apart, with the first delivery occurring at 7s from cue onset. CS3 was a visual stimulus (10s flashing cue light, 1s on, 2s off) and had no consequence (CS-). There were 8 presentations of each cue (inter-trial interval: 120s VI) presented in pseudorandomised order so that no more than two of the same cue was presented in sequence. All animals received the same order of cue presentation. A photocell beam detected magazine entries.

#### Reward omission test 1–extinction

Following appetitive conditioning, rats received a single test session where CS2 was extinguished while CS1 and CS3 were maintained as CS+ and CS-, respectively (**Figure 1A**).

**Figure 1.**
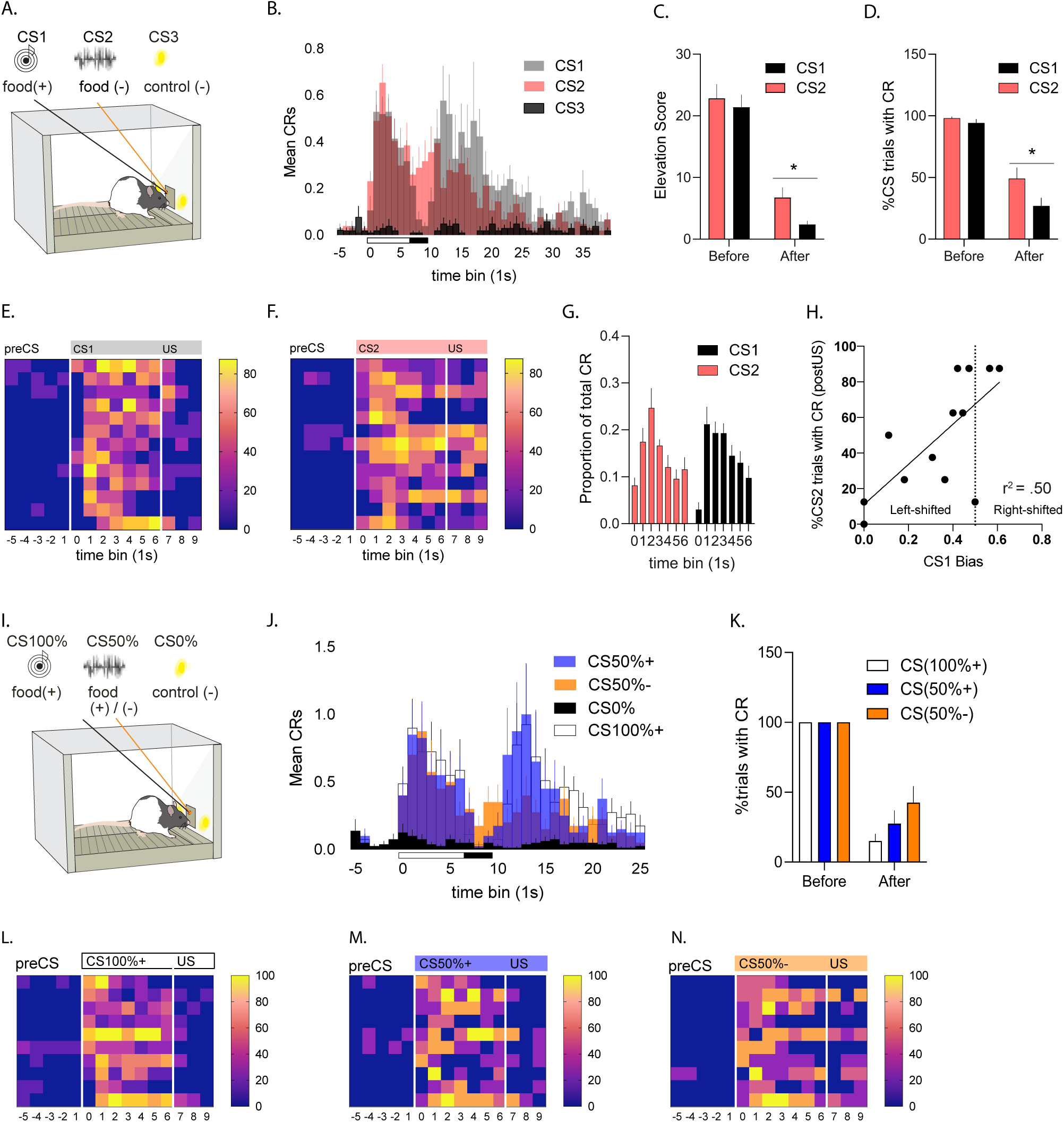
Characterization of dynamic appetitive shifts in food-seeking behavior. ***A.*** Schematic depicting reward omission test–extinction. CS1: food-reinforced; CS2: unexpected food omission; CS3: never reinforced. ***B.*** Average seeking responses per 1s time bin across CS1, CS2, and CS3. ***C.*** Mean CS-generated seeking responses before and after US delivery for CS1 and equivalent time for CS2; elevation relative to CS3. ***D.*** Percentage of trials paired with seeking before and after US delivery for CS1 and equivalent time for CS2. ***E-F.*** Heatplot depicting percentage of trials paired with seeking per 1s time bin (columns) per subject (rows) for CS1 (***E.***) and CS2 (***F.***). ***G.*** Shifts in seeking response following CS onset prior to US delivery (CS1) or equivalent time (CS2). ***H.*** Correlation of response bias for CS1 with percentage of trials paired with persistent seeking for CS2. ***I.*** Schematic depicting partial reward omission test. CS100%: always food-reinforced; CS50%: partially food-reinforced; CS3: never reinforced. ***J.*** Average seeking responses per 1s time bin across conditions: CS100%+, CS50%+, CS50%- and CS0%. ***K.*** Percentage of trials paired with seeking before and after US delivery for CS100% and CS50% and equivalent time for CS50%-. ***L-N.*** Heatplot depicting percentage of trials paired with seeking per 1s time bin (columns) per subject (rows) for CS100%+ (***L.***), CS50%+ (***M.***), and CS50%- (***N.***). *\*p<.05*.

#### Reward omission test 2–partial reinforcement

In a separate cohort, following appetitive conditioning, rats received three sessions where CS2 was partially reinforced (CS50%) while CS1 and CS3 were maintained as CS+ (CS100%) and CS-(CS0%) respectively (**Figure 1I**). Food was paired with CS50% on alternating trials, starting at trial 1 for half the animals, and on trial 2 for the remaining animals.

#### Optogenetic silencing for BLA→AcbSh and BLA→AcbC pathways

Following appetitive conditioning, CS2 was extinguished while CS1 and CS3 were maintained as CS+ and CS-, respectively. Responding to CS2 was reduced to criterion (CS1: CS2 ratio >.6 over two consecutive days, approximately 4-7 days). Rats were subsequently tested under photostimulation of eNpHR3.0 to silence the BLA→AcbSh or BLA→AcbC pathways. CS presentations were pseudorandomised with the restriction that each CS type was presented in the first three test trials, and again in the next three trials. On the following day, rats were tested again under the same behavioral parameters but in the absence of photostimulation (post-test).

For light delivery, an LED (625nm wavelength, Doric Instruments) was suspended to a counterweighted arm above the chamber. It was connected to two custom patch cables (400µm ⌀) that mated with bilaterally implanted optic fiber ferrules on the rat. Photostimulation (constant pulse, ∼7-10mW illumination at fiber tip) were triggered via an LED driver (Doric Lenses, Quebec, Canada) 200ms prior to each CS onset. For tests involving across-trial silencing, photostimulation were 20s in duration and therefore extended across the entire length of the CS as well as a 10s post CS period. For tests involving within-trial silencing (i.e., before or after scheduled US), photostimulation were 7s in duration. Half the rats were tested so that the first three CSs were paired with photostimulation, followed by every other trial thereafter. The remainder received the inverse pattern. Thus, photostimulation was delivered intermittently across trials so that for each CS type, 50% of trials were ‘On’ and 50% were ‘Off’ conditions.

#### Brain Slice Preparation and Electrophysiological recordings

Brain slices were prepared > 4 weeks after viral injections according to standard procedures. Rats were deeply anesthetized with isoflurane (5%) and killed via decapitation. Brains were extracted and immersed in ice-cold sucrose-modified artificial cerebral spinal fluid (ACSF), with the following composition (in mM): sucrose, 124; NaCl, 62.5; KCl, 2.5; NaHCO3, 26; NaH2PO4, 1.2; glucose, 10; CaCl2, 0.5; and MgCl2, 3.3 (equilibrated with 95% CO2, 5% O2). Coronal brain slices (300 µm) were prepared using a VT1200 vibratome (Leica, Wetzlar, Germany). Slices were then transferred to a holding chamber containing standard ACSF (119 mM NaCl, 2.5 mM KCl, 1.3 mM MgCl2, 2.5 mM CaCl2, 1.0 mM Na2H2PO4, 26.2 mM NaHCO3, 11 mM glucose, equilibrated with 95% CO2, 5% O2) and incubated at 32 °C for 10 minutes in Braincubator (Payo Scientific, Sydney, Australia) and then maintained at 16 °C until recording.

For electrophysiological recordings, slices were transferred to a recording chamber and perfused (2 ml min-1) with ACSF (30 °C). Cells were visualized using a Zeiss Axio Examiner D1 microscope equipped with a 20x water immersion objective (1.0 NA), an LED fluorescence illumination system (PE2, CoolLED, Andover, UK), an electron multiplying charge coupled device camera (iXon, Andor). Targeted whole-cell patch-clamp recordings were made from red-fluorescent cells in the AcbSh using patch pipettes (2-5 MΩ) filled with an internal solution containing (in mM) 135 KMeSO4, 8 NaCl, 10 HEPES, 2 Mg2-ATP, 0.3 Na3-GTP, 0.3 EGTA, 0.05 Alexa 594 (pH 7.3 with KOH, 280-290 mOsm). Electrophysiological recordings were amplified and filtered using a Multiclamp 700B amplifier (Molecular Devices, California, USA) digitized at 20 kHz (Digidata1440A, Molecular Devices), and controlled and analyzed using AxoGraph (Axograph, Sydney, Australia). Only cells with action potential amplitudes >90 mV, and input resistances > 60 MΩ were considered healthy and included in the data set. ChR2 infected terminals were stimulated using brief (1-10 ms) 470 nm LED illumination delivered through the objective. Neurons were considered connected if the light reliably evoked a synaptic current > 15 pA (Vm = - 70 mV). Light-evoked responses and latency of light-evoked responses were calculated from light onset.

#### Immunohistochemistry and Histology

At the conclusion of the optogenetic and retrograde tracing studies, rats were deeply anesthetized with sodium pentobarbital (100mg/kg, i.p.) and perfused transcardially with 50ml of 0.9% saline followed by 400ml of 4% paraformaldehyde in 0.1M phosphate buffer (PB), pH7.4. Brains were postfixed for 1h in the same fixative and placed in 20% sucrose overnight. For optogenetic studies, tissue were sectioned coronally at 40µm using a cryostat and stored in a phosphate buffered solution (PBS). Sections were processed using fluorescent immunohistochemistry for detection of eYFP. Free floating sections were washed in a solution of PBS with 0.2% Triton X-100 and 0.2% bovine serum albumin (PBST) for 20min at room temperature and then incubated in a blocking solution made of PBST with 10% normal donkey serum (Jackson ImmunoResearch, 017-000-121) for 30 min. Sections were incubated overnight at 4°C with gentle agitation in a chicken antiserum against GFP (1:1500, Invitrogen, A11120). After subsequent 3 x 10 min washes in PBST, sections were incubated in a secondary antibody (AlexaFluor 488 donkey anti-chicken, 1:400, Invitrogen, A32766) in PBST for 2h at room temperature. Finally, sections were washed in PBS, mounted and coverslipped with PermaFluor Aqueous Mounting Medium (ThermoFisher Scientific, TA030FM). Images were digitally captured using a (Hamamatsu ORCA-Flash4.0) attached to an Olympus BX53 epifluorescent microscope and CellSens software. For the CTb-tract tracing study, perfused and post-fixed tissue were sectioned at 35µm, mounted and coverslipped. For cell counting, mounted amygdala sections (140µm apart) were imaged as described above and both single- (red or green) and double-labeled (yellow) cells were counted manually across 8 sections. Anterior amygdala counts were derived from the first 4 sections, posterior amygdala counts from the remaining sections.

#### Experimental Design and Statistical Analysis

The experiments used single factor (CS type) or (3) (CS Type) x 2 (eYFP, eNpHR3.0) x (2) (light On vs Off) factorial designs. The primary data of interest were magazine entries following the CS, including periods before (0-7s) and after (7-10s) the US. These were automatically collected via the MedAssociates software. These data were analyzed using a mixed-model multivariate ANOVA with between-subjects (group) and within-subjects (CS Type, photoinhibition on vs off) factors. Significance was assessed against a type I error rate of 0.05.

## Results

### Reward produces a pause in appetitive seeking

High levels of seeking during foraging, pursuit, or predation, can quickly transition into behavioral quiescence during reward receipt and consumption. An account for this transition is that reward receipt demands different behavioral responses, and shunting seeking-generated activity is necessary to facilitate this behavioral switch. So, we first asked whether reward receipt is indeed a critical driver of reductions in seeking.

We used a Pavlovian conditioning preparation to study changes in appetitive seeking (magazine approach) around periods of consumption (N=13). We measured seeking as stimulus-signaled magazine entries to three distinct stimuli. Conditioned stimulus (CS1) was an established CS+ paired with food reward, CS2 was formerly a CS+ but reward was withheld, and CS3 was never paired with reward (CS-) (**Figure 1A**). Thus, CS1 required an appetitive-consummatory transition around the time of reward delivery; CS2 provided a measure of appetitive change during reward omission; and CS3 was neutral. Under these conditions, the expression of seeking and consumption were temporally segregated to the onset of the CS+ and US, respectively.

The profile of seeking varied across CS types (**Figure 1B**). Seeking responses to CS1 were bimodally distributed. Mean levels of seeking peaked in the first 3 seconds from cue onset, then declined by the end of CS1, coinciding closely with delivery of the reward (food) US. During the reward delivery period, new responses to CS1 were nil in 73% of observation trials. At the offset of CS1, seeking subsequently recovered with some variability, then declined at a protracted pace. This trend reflects a robust seeking response to CS1 that is significantly and transiently reduced after reward delivery (CS- vs US-generated response means across 1s timebins, F_(1,12)_=36.021, p<.001; US- vs postCS-generated response means across 1s timebins, F_(1,12)_=5.597, p=.036; **Figure 1B**). We observed a similar reduction in seeking to the non-reinforced appetitive cue (CS2) in the period of expected reward delivery (CS- vs US-generated response means across 1s timebins, F_(1,12)_=7.814, p<.05, **Figure 1B**) although this reduction was significantly less than during CS1 (2-way CS type x appetitive phase interaction: F_(1,12)_ = 6.144, p=.029; simple effects on consumption phase, CS1 v CS2: F_(1,12)_ = 6.516, p=.025, **Figure 1B**). In contrast to CS1 and CS2, performance to the never conditioned stimulus, CS3, was low throughout this period (**Figure 1B**).

To confirm that the reduction of seeking to CS1 was due to the presence of reward rather than other factors (e.g., fatigue) we compared levels of seeking during CS1 and CS2 in the periods before and after expected reward delivery (**Figure 1C**). Critically, levels of seeking significantly varied between the cues after (F_(1,12)_ = 9.399, *p*=.010, Figure 1C) but not before reward delivery (F_(1,12)_ = .573, *p*=.464, **Figure 1C**). We also compared between the two cues the proportion of trials that elicited seeking both before and after reward delivery. CS2 elicited seeking on a significantly greater proportion of trials than CS1 after reward delivery (F_(1,12)_ = 7.149, p=.020) but not before (F_(1,12)_ = 1.157, p=.303) (**Figures 1D-F**). So, omission of an expected reward (ie, CS2) promotes persistence of seeking, suggesting that reward delivery is sufficient to transiently suppress this form of appetitive behavior.

To confirm that these conclusions were not specific to comparing CS1 to the extinguished CS2, we studied how seeking changed under partial reinforcement. Separate animals (N=10) were trained to three distinct stimuli, where CS100% always predicted reward (100% reward), CS50% only partially predicted reward (50% reward), and CS0% never predicted reward (0% reward) (**Figure 1I**). The behavioral dynamics of seeking were similarly bimodal as before (**Figure 1J**). Critically, seeking was significantly reduced following reward delivery (CS vs US-generated response means across 1s timebins, CS100%+: F_(1,9)_=18.054, p=.002; CS0%+: F_(1,9)_=24.125, p=.001; **Figure 1J**) however there were no significant differences between US- and postCS seeking activity (1s timebins, CS100%+: F_(1,9)_=3.919; p=.079; CS50%+: F_(1,9)_=3.77, p=.084; **Figure 1J**) largely owing to variability in the post-CS period. As before, performance to the never conditioned stimulus, CS0%, remained at baseline throughout (**Figure 1J**).

Closer inspection of behavior elicited by the partially-reinforced CS50% showed a significant reduction in seeking around the time of expected reward delivery on trials that ended in reward omission (CS50%-; before vs after expected reward response means across 1s timebins: F_(1,9)_=45.489, p<.001; **Figure 1J**) however as observed previously, this reduction in performance was significantly smaller than that during CS100%+ (2-way CS type x CS component (before v after reward) interaction: F_(1,9)_ = 6.815, p=.028; simple effects on responding after reward, CS100%+ vs CS50%-: F_(1,9)_ = 7.477, p=.023, **Figure 1J**). This reduction was also smaller relative to reward reinforced trials with the same cue, CS50%+ (CS50%+; 2-way CS type x CS component interaction: F_(1,9)_ = 7.732, p=.021; simple effects on seeking after reward delivery, CS50%+ vs CS50%-: F_(1,9)_ = 4.808, p=.056, **Figure 1J**). This confirms that reward delivery promotes suppression of seeking.

Consistent with this, the proportion of trials eliciting a seeking response was significantly higher for the nonreinforced CS50%-than for the reinforced CS100%+, after reward delivery (2 way CS type x CS component interaction: F_(1,9)_ = 10.471, p=.004; simple effects on responding ‘after reward’, CS100%+ vs CS50%-: F_(1,9)_ = 10.471, p=.004, **Figures 1K-N**). However performance generated by CS50% on reinforced vs non reinforced trials were not significantly different on this measure (2 way CS type x CS component interaction: F_(1,9)_ = 1.976, p=.194, **Figure 1K-N**).

Together, these findings show that the presence of the appetitive CS robustly generates seeking responses, which in turn are rapidly suppressed following reward delivery. Moreover, the presence of the reward is a primary driver of this transition.

### Appetitive restraint occurs prior to reward delivery

Reward was not the only driver of changes in seeking. Although seeking peaked prior to reward delivery, the timing of this peak was premature relative to the precise time of reward delivery (**Figure 1B**). So, we inspected this trend more closely.

The distribution of CS-generated seeking before reward delivery increased as a function of time, and then decreased immediately prior to reward delivery (Experiment 1: quadratic contrast for CS1, magazine entries as a proportion of preUS CR, F_(1,12)_ = 35.802, p=.000; and CS2: F(1,12) = 12.733, p=.004, **Figure 1G**). We next asked whether this inflection in seeking was established prior to test, or whether it was an artifact of the testing conditions, given that cue-reward contingencies were changed on test for CS2. For simplicity, we addressed this by sectioning the pre-reward period (from CS onset to US onset) into three time intervals relative to reward delivery: 1^st^ (0-2.33sec), 2^nd^ (2.33-4.66sec), and 3^rd^ (4.66-7sec) timebin on each day of training and on test. A significant left-shifted peak in seeking activity emerged on the last two days of training (quadratic contrast for Day 7 CS1: F_(1,12)_ = 12.527, p=.004; Day 7 CS2: F_(1,12)_ = 6.326, p=.027; Day 8 CS1: F_(1,12)_ = 8.459, p=.013; Day 8 CS1: F_(1,12)_ = 4.533, p=.055) indicating that the suppression of seeking before reward delivery was not an artifact of test.

We considered the possibility that this marked behavioral change, expressed in the seconds prior to reward delivery, may be necessary for the subsequent transition from seeking to consumption. However, although the downshift emerged at the end of training, significant seeking-taking inflections were detected much earlier in training (portion of trials with CR at CS vs US onset: Day3 CS1 F_(1,12)_ = 8.766, p=.012; CS2 p>0.05; Day 4 CS1 F_(1,12)_ = 4.868, p=.048; CS2 F_(1,12)_ = 21.056, p=0.001). This suggests that the downshift in appetitive behavior prior to reward delivery was not required for suppression of appetitive seeking during reward delivery.

We then explored whether the appetitive restraint was related to the behavioral inflection around reward delivery. To address this, we measured individual seeking bias towards or away from the time of reward delivery using bias scores relative to the time periods intermediate and proximal to reward onset (2^nd^ and 3^rd^ timebins, intervals defined previously). We calculated right-shifted bias scores for CS1 as the average proportion of pre-reward CS responses within the 3^rd^ timebin / sum of average proportion of pre-reward CS responses during the 2^nd^ and 3^rd^ time bins (i.e., scores >.5 indicate that the subject’s response preference was right-shifted, biased towards the 3^rd^ timebin). The majority of animals showed a left-shifted preference for at least one of the cues (bias <.5 in 11 of 13 rats, **Figure 1H**). Critically, response bias relative to reward delivery was not significantly associated with pauses in seeking following reward delivery on test (CS1 bias, CS1 proportion of trials with postUS CR, F_(1,11)_=2.623, p=.1136). Instead, it was significantly associated with levels of seeking when reward was absent (CS1 bias, CS2 proportion of trials with postUS CR, R^2^=.500; F_(1,11)_ = 10.99, p=.0069, **Figure 1H**), and thus related to the strength or persistence of the CS-generated response.

Together these findings demonstrate appetitive restraint in the period prior to reward delivery, but this restraint is not obviously related to the pauses driven by reward delivery. Rather, it is closely associated with motivational changes occurring within the trial. So, appetitive seeking is a sculpted response with two points of transition: one after reward delivery due to presence of the reward and one before reward delivery due to dynamic motivational fluctuations.

### BLA→AcbSh pathway mediates appetitive transitions

The nucleus accumbens (Acb), and its major glutamatergic input from the basolateral amygdala (BLA), may play a crucial role in these appetitive-consummatory transitions. The Acb comprises shell and core regions, so we first determined segregation among the BLA→AcbSh and BLA→AcbC pathways. We applied retrograde tracer (CTb555 and CTb488) to both AcbSh and AcbC so that single and dual-labeled (i.e. collateral AcbSh+AcbC targeting) BLA projections would be indicated (N=4). Injections for each pair were restricted to the targeted area and were of similar size in spread (example in **Figure 2B**). The injections were largely confined to the dorsomedial AcbSh and to the laterally adjacent region including AcbC (**Figures 2A-B**), which were targets of our subsequent studies. AcbSh and AcbC projector cells in the amygdala were counted in the basal amygdala (BA) and adjacent lateral amygdala (LA) regions, across anterior and posterior segments. **Figure 2C** shows the distribution of mean counts of labelled cells in BLA along its anterior-posterior axis and also between BA and LA nuclei. There were no significant differences in the total numbers of AcbSh projectors and AcbC projectors (**Figure 2D**). However, there were some distinctions.

**Figure 2.**
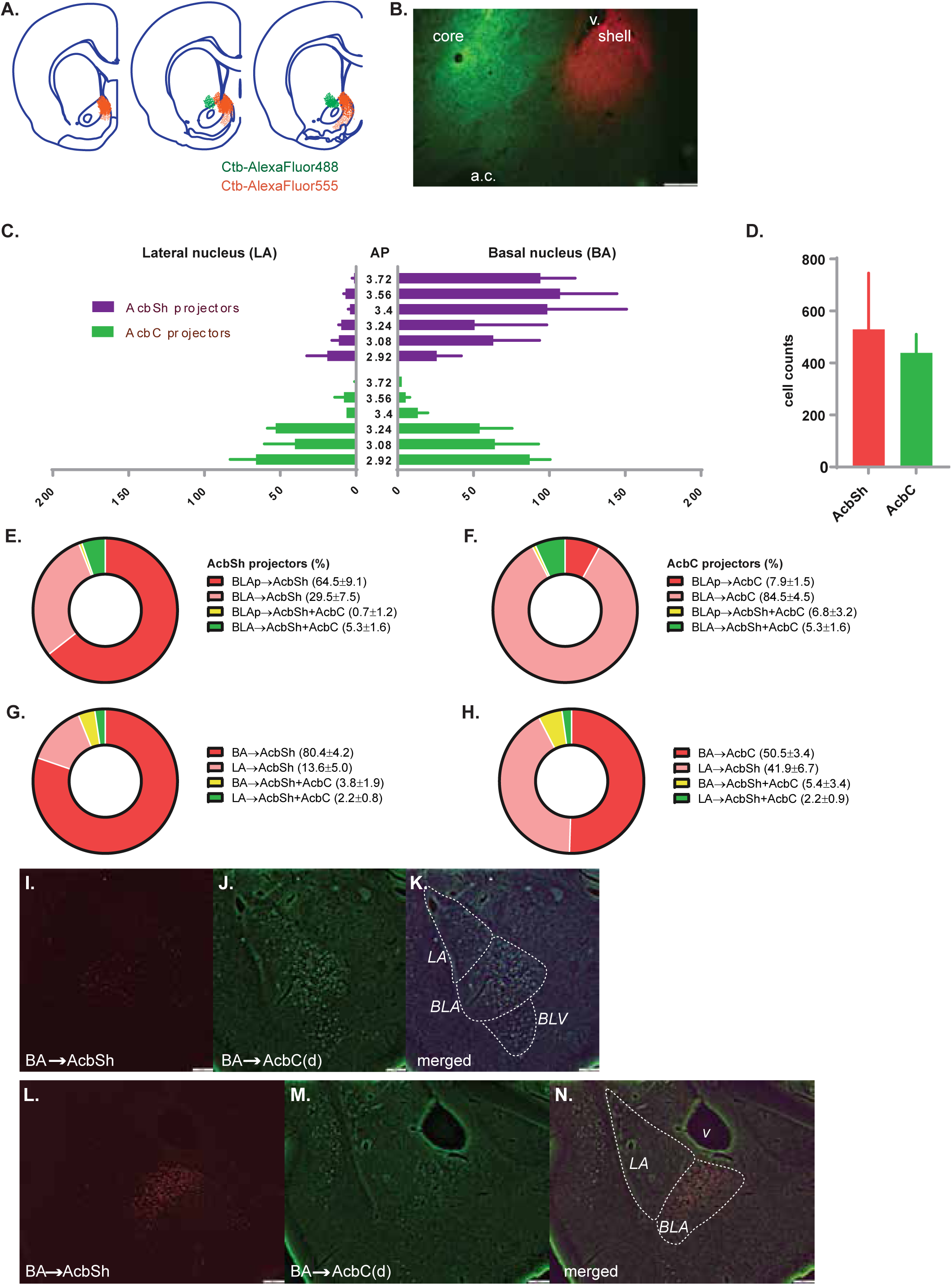
Characterization of BLA projectors to AcbSh and AcbC. ***A.*** Location of CTb-AlexaFluor488 and CTb-AlexaFluor555 deposits. ***B.*** Photomicrograph of representative dual-tracer deposits in AcbSh and AcbC. ***C.*** Topographical distribution of AcbSh- and AcbC-projectors across the rostro-caudal axis of the amygdala for basal (BA) and lateral (LA) subregions. ***D.*** Frequency of AcbSh- and AcbC-projectors counted in amygdala. ***E.*** Percentage of AcbSh-projectors in BLA localized within posterior (red = single labeled, yellow = dual-labeled) and anterior (pink = single labeled, green = dual labeled) BLA. ***F.*** Percentage of AcbC-projectors in BLA localized within posterior (red = single labeled, yellow = dual-labeled) and anterior (pink = single labeled, green = dual labeled) BLA. ***G.*** Percentage of AcbSh-projectors in amygdala localized within BA (red = single labeled, yellow = dual-labeled) and LA (pink = single labeled, green = dual labeled) BLA. ***H.*** Percentage of AcbC-projectors in amygdala localized within BA (red = single labeled, yellow = dual-labeled) and LA (pink = single labeled, green = dual labeled) BLA. ***I-K:*** Representative photomicrographs of anterior BLA. Few AcbSh-projectors labeled in ***I.*** AcbC-projectors in ***J.*** Merged image in ***K. L-N:*** Representative photomicrographs of posterior BLA. AcbSh-projectors labeled in ***L.*** AcbC-projectors in ***M.*** Merged image in ***N*.**

Among AcbSh-projecting BLA neurons, a majority were localized to the posterior BLA (BLAp), which included the lateral (BLP) and medial (BMP) nuclei of the BA (BLAp: 64.46%±9.07 vs BLAa: 29.50%±7.50; **Figure 2E**), and only few cells were dual-labelled (BLAp: 0.71%±.27 vs BLAa: 5.33%±1.6; **Figure 2E**), indicating largely segregated AcbSh projector neurons from AcbC projectors in the BLA. Consistent with this, we found that among AcbC-projecting BLA neurons, a majority were localized to the anterior BLA (BLAa: 29.50%±7.50 vs BLAp: 7.93%±1.45; **Figure 2F**), and similar to AcbSh projectors, there were few dual-labelled cells (BLAp: 0.74%±.32 vs BLAa: 6.8%±3.2; **Figure 2F**). These findings indicate largely non-overlapping BLA→AcbSh and BLA→AcbC neurons that intermingle moderately in the anterior regions of the BLA. They support previous reports of a topographical anterior-posterior organization of BLA neurons (Wright, Beijer, & Groenewegen, 1996; Zahm & Brog, 1992) targeting the ventral striatum.

We also found segregation among AcbSh and AcbC projectors within BLA subnuclei. Specifically, for AcbSh projectors, a majority were non-overlapping and localized primarily within the BA (BA: 80.37±4.20 vs LA: 13.59±4.95; dual-labelled in BA: 3.80±1.90 vs LA: 2.24±0.85; **Figure 2G**). Similarly, for AcbC projectors, a majority were non-overlapping, and about half were localized to the BA and the other in the LA (BA: 50.49±3.40 vs LA: 41.93±6.70; dual-labelled in BA: 5.39±3.37 vs LA: 2.20±0.94; **Figure 2H**). Together these findings suggest largely segregated neurons form the BLA→AcbSh and BLA →AcbC pathways, with segregation organized along the anterior-posterior axis of the BLA as well between BA and LA nuclei.

We next used an optogenetic approach to selectively silence the BLA→AcbSh (eYFP n = 6, eNpHR3.0 n = 7) and BLA→AcbC pathways and determined their effects on the appetitive transitions that occur after and before reward delivery. We again studied seeking in relation to three distinct stimuli, where CS1 was an established CS+, CS2 was formerly a CS+ but was subsequently extinguished, and CS3 was a CS- (**Figure 3E**).

**Figure 3.**
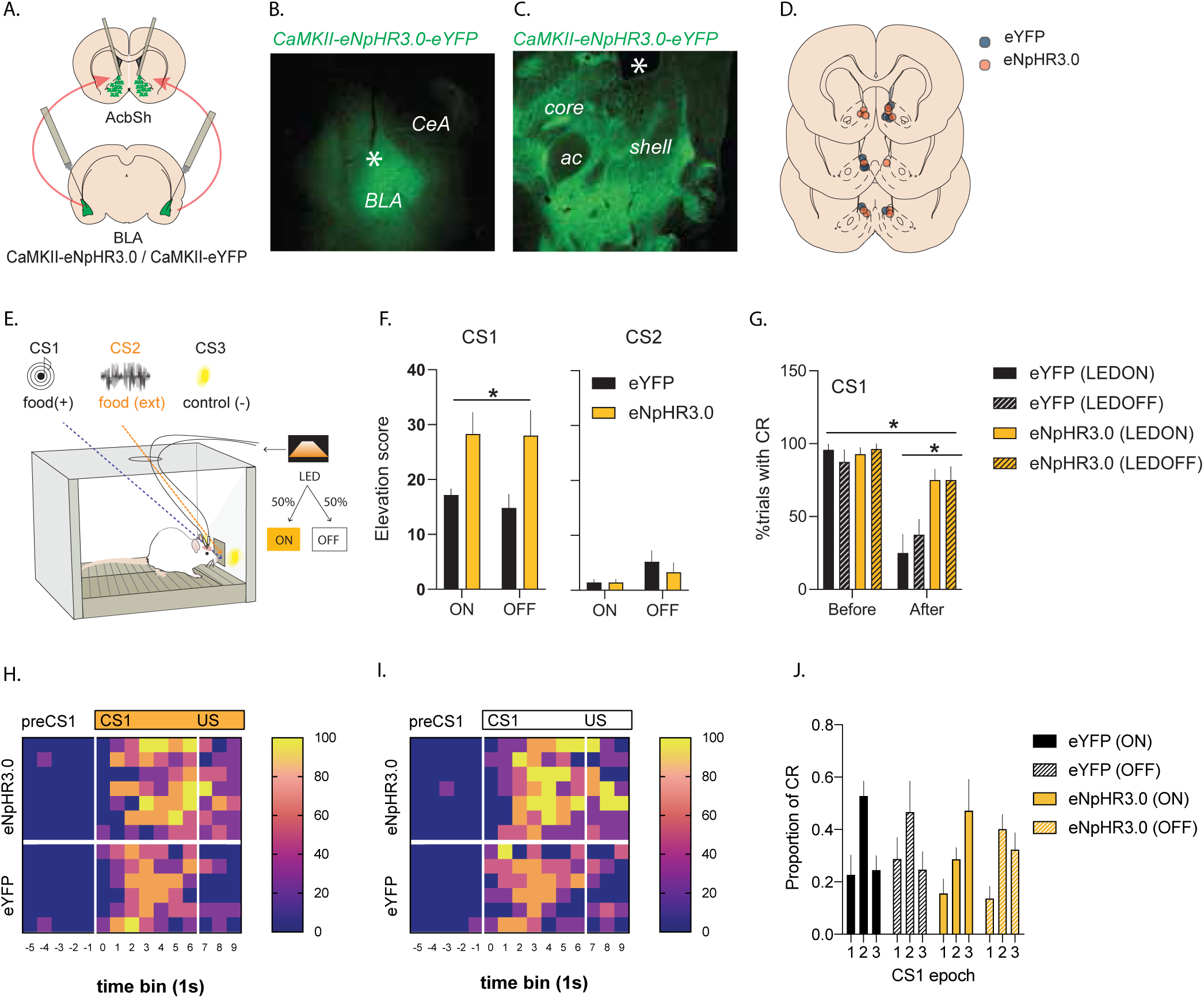
Optogenetic silencing of BLA→AcbSh on appetitive transitions. ***A.*** Schematic of eYFP or eNpHR3.0 injected in BLA and fiber optic cannulae positioned above AcbSh. ***B.*** Representative photomicrograph of spread of eNpHR3.0-eYFP in BLA, red arrow = injection tip. ***C.*** Representative photomicrograph of BLA terminal labeling in accumbens, red arrow = cannula tip above dorsal AcbSh. ***D.*** Locations of fiber optic cannulae in AcbSh mapped on coronal sections. ***E.*** Schematic depicting appetitive test. CS1: food-reinforced; CS2: extinguished; CS3: never reinforced; LED delivered on 50% of trials of each CS. ***F.*** Mean CS-generated seeking responses for CS1 and CS2. ***G.*** Percentage of trials paired with seeking before and after US delivery for CS1. ***H-I.*** Heatplot depicting percentage of trials paired with seeking per 1s time bin (columns) per subject (rows) for CS1 with (***H.***) and without (***I.***) photoinhibition. ***J.*** Shifts in seeking response following CS1 onset prior to US delivery.

Silencing BLA→AcbSh during the entire CS+ (i.e. CS and US delivery) significantly increased overall seeking (CS1 group main effect: eNpHR3.0 vs eYFP, F_(1,11)_= 7.44, *p=*.02; **Figure 3F**) and this effect persisted across CS+ trials, including on trials without photoinhibition (interaction group x light (On vs Off): F_(1,11)_=.192, *p*=.67, **Figure 3F**). To rule out non-specific differences between groups, we confirmed that these effects were selective to the CS+. There were no differences between groups in responding to the extinguished CS (CS2, group main effect: F_(1,11)_=.32, *p*=.583; interaction group x light: F_(1,11)_=.569, *p*=.466; **Figure 3F**) or to the CS- (CS3, group main effect: F_(1,11)_=.339, *p*=.572; interaction group x light: F_(1,11)_=.451, *p*=.516). The effects were also selective to the test conditions because there were no differences between groups on the following day (CS1 group main effect: F_(1,11)_=.03, *p=*.866; group x light interaction: F_(1,11)_=1.402, *p=*.261). These findings suggest that silencing BLA→AcbSh inflates seeking and, given its specificity to the CS+, that this inflation is not due to a non-selective change in baseline performance or to disinhibition of non-rewarded seeking. Thus, we next asked whether this inflation was driven, at least in part, by disruptions to transitions following reward delivery.

To do this, we inspected behavior in the periods before and after reward delivery. As expected, the proportion of trials that elicited a seeking response was significantly greater before than after reward (main effect: before vs. after, F_1,11_=23.307, p=.001, **Figure 3G**) indicating a significant pause in seeking following reward delivery (**Figures 3G-I**). Critically, silencing BLA→AcbSh pathway significantly impaired this suppression (group x CS phase, F_1,11_=6.045, p=.032; CS phase (after reward) simple effects for group, LED On: F_1,11_=11.846, p=.006; and group, LED Off: F_1,11_=6.953, p=.023, **Figure 3G**). This suggests that reward presence controls seeking via a BLA→AcbSh pathway.

We also asked whether BLA→AcbSh pathway contributed to transitions that occurred prior to reward delivery. First, we confirmed that the distribution of seeking before reward was overall significantly quadratic, indicating that magazine entries increased as a function of time, and then decreased immediately prior to reward delivery (quadratic contrast for proportion of preUS CR per epoch across groups, F_(1,11)_ = 9.766, p=.010, **Figure 3J**). Critically, there was a significant group x quadratic trend interaction for the light on condition (F_(1,11)_ = 7.790, p=.018) but not the light off condition (F_(1,11)_ = .053, p=.822) suggesting that the profile of seeking during trials involving photoinhibition was significantly different between groups. Indeed, on closer inspection, seeking was significantly quadratic in profile in the control group, eYFP, (quadratic trend, On condition: F_(1,5)_=11.333, p=.02) but not in the eNpHR3.0 group (quadratic trend, On condition: F_(1,6)_=.050, p=.830, **Figure 3J**). Instead, the distribution of seeking for group eNpHR3.0 followed a significant linear profile (On condition: F_(1,6)_=6.410, p=.045; Off condition: F_(1,6)_=6.075, p=.049, **Figure 3J**) indicating that seeking in this group peaked in the period most proximal to reward delivery. This suggests that silencing BLA→AcbSh pathway prevented suppression of seeking prior to reward delivery. Moreover, this effect was not simply due to an elevation of the appetitive response given that responses on this measure were normalized as proportions of total seeking prior to reward and since group differences in frequency of seeking before reward were not significant (magazine entries, group main effect, F_(1,11)=_4.211, p=.065). Rather, the findings suggest that silencing BLA→AcbSh pathway modified the topography of the cue-generated appetitive response.

We applied the same approach to study the contribution of BLA→AcbC pathway (eYFP n = 6, eNpHR3.0 = 8; **Figures 4A-D**). There was no significant difference in seeking between groups to the CS+ (CS1 group main effect, F_(1,12)_=.314, p=.586; group x light, F_(1,12)_=.177, p=.681, **Figure 4F**), to the extinguished CS (CS2 group main effect, F_(1,12)_=.314, p=.586; group x light, F_(1,12)_=.177, p=.681, **Figure 4F**) or the non-reinforced CS (CS3 group main effect, F_(1,12)_=.62, p=.446; group x light, F_(1,12)_=1.302, p=.276) on test. As expected, the proportion of trials that elicited a seeking response was significantly higher before than after reward delivery (main effect: appetitive vs. consummatory, F_1,12_=41.787, p=.000, **Figure 4G-I**). This confirms the suppression of seeking behavior by reward delivery. Critically, and in contrast to the BLA → AcbSh, silencing BLA→AcbC pathway did not affect this suppression (consummatory period: group x light, F_1,12_=1.886, p=.195; group, LED On, F_1,12_=3.231; p=.097; group, LED Off: F_1,12_=.135, p=.72, **Figure 4G**). However, it did have a significant effect on the vigor (seeking/second) of seeking responses after reward delivery in a light-dependent manner (consummatory period: group x light, F(1,12)=4.994, p=.045; simple effects group, On, F_(1,12)_ = 5.675, p=.035; Off, F_(1,12)_ =.164, p=.693). There was no effect of BLA→AcbC silencing on the vigor of active seeking prior to reward delivery (appetitive period: group x light, F(1,12)=.606, p=.045; simple effects group, On, F_(1,12)_ = .827, p=.381; Off, F_(1,12)_ =.013, p=.911).

**Figure 4.**
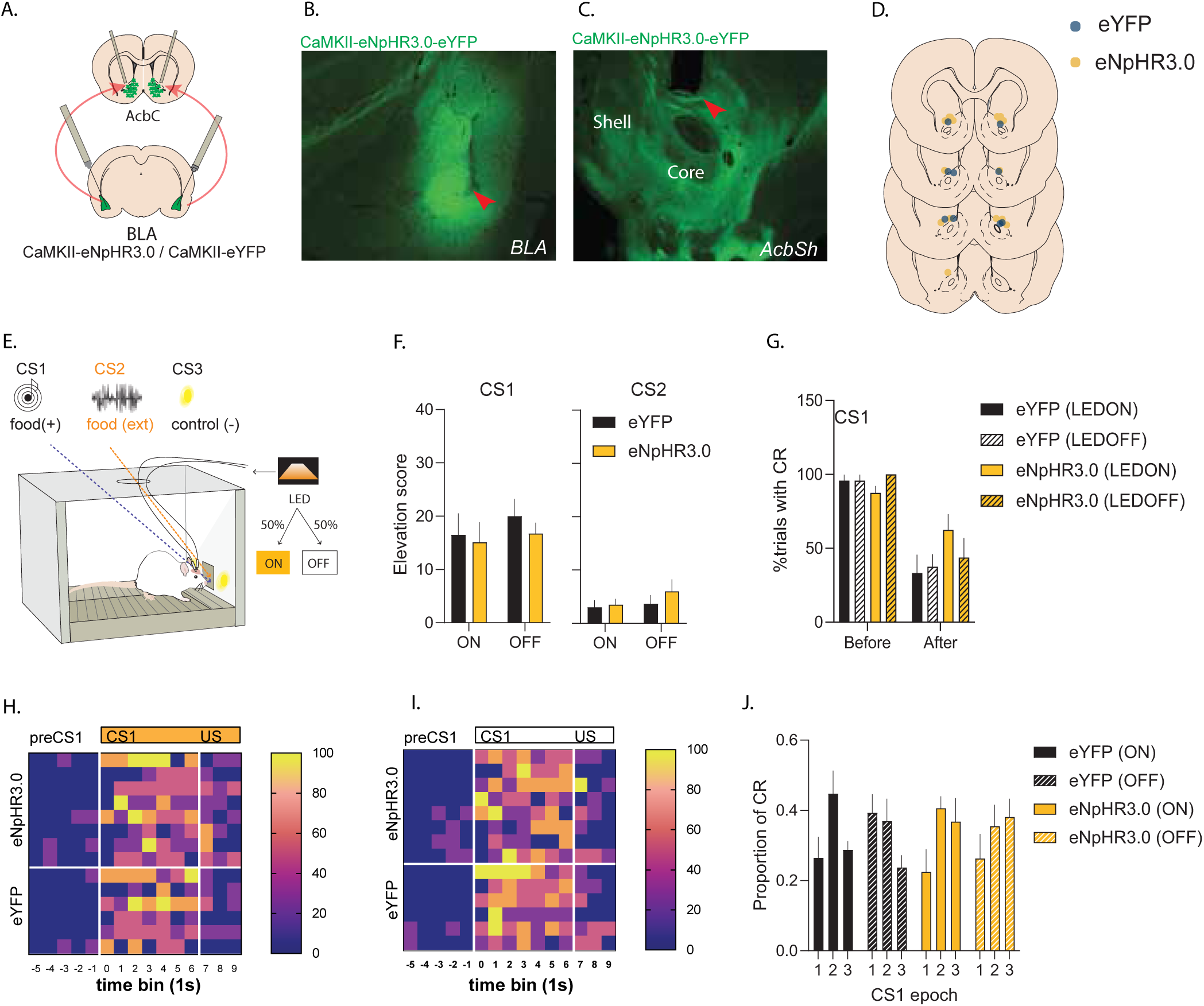
Optogenetic silencing of BLA→AcbC on appetitive transitions. ***A.*** Schematic of eYFP or eNpHR3.0 injected in BLA and fiber optic cannulae positioned above AcbC. ***B.*** Representative photomicrograph of spread of eNpHR3.0-eYFP in BLA, red arrow = injection tip. ***C.*** Representative photomicrograph of BLA terminal labeling in accumbens, red arrow = cannula tip above AcbC. ***D.*** Locations of fiber optic cannulae in AcbC mapped on coronal sections. ***E.*** Schematic depicting appetitive test. CS1: food-reinforced; CS2: extinguished; CS3: never reinforced; LED delivered on 50% of trials of each CS. ***F.*** Mean CS-generated seeking responses for CS1 and CS2. ***G.*** Percentage of trials paired with seeking before and after US delivery for CS1. ***H-I.*** Heatplot depicting percentage of trials paired with seeking per 1s time bin (columns) per subject (rows) for CS1 with (***H.***) and without (***I.***) photoinhibition. ***J.*** Shifts in seeking response following CS1 onset prior to US delivery.

We then asked whether BLA→AcbC pathway was required for transitions that occurred prior to reward delivery. It was not. While appetitive responding increased and then decreased prior to reward delivery (quadratic contrast for preUS CR, F_(1,12)_ =4.951, p=.046, Figure 4J), there were no significant differences between groups (group x quadratic trend interaction: F_(1,12)_ =.251, p=.625; light on: F_(1,12)_ =.364, p=.558; light off: F_(1,12)_ =.024, p=.879), **Figure 4J**.

### Silencing BLA→AcbSh pathway reveals interdependence of appetitive transitions

Silencing the BLA→AcbSh pathway augmented seeking behavior during periods of appetitive restraint: immediately prior to, and following, reward delivery. So, we asked if these effects were interdependent or domain specific. To test this, we silenced the BLA→AcbSh pathway (eYFP n = 10, eNpHR3.0 = 8) in the period after (**Figure 5E**) or before (**Figure 5L)** reward delivery to target these transitions separately, and asked, does silencing during one transition affect behavior during the other?

**Figure 5.**
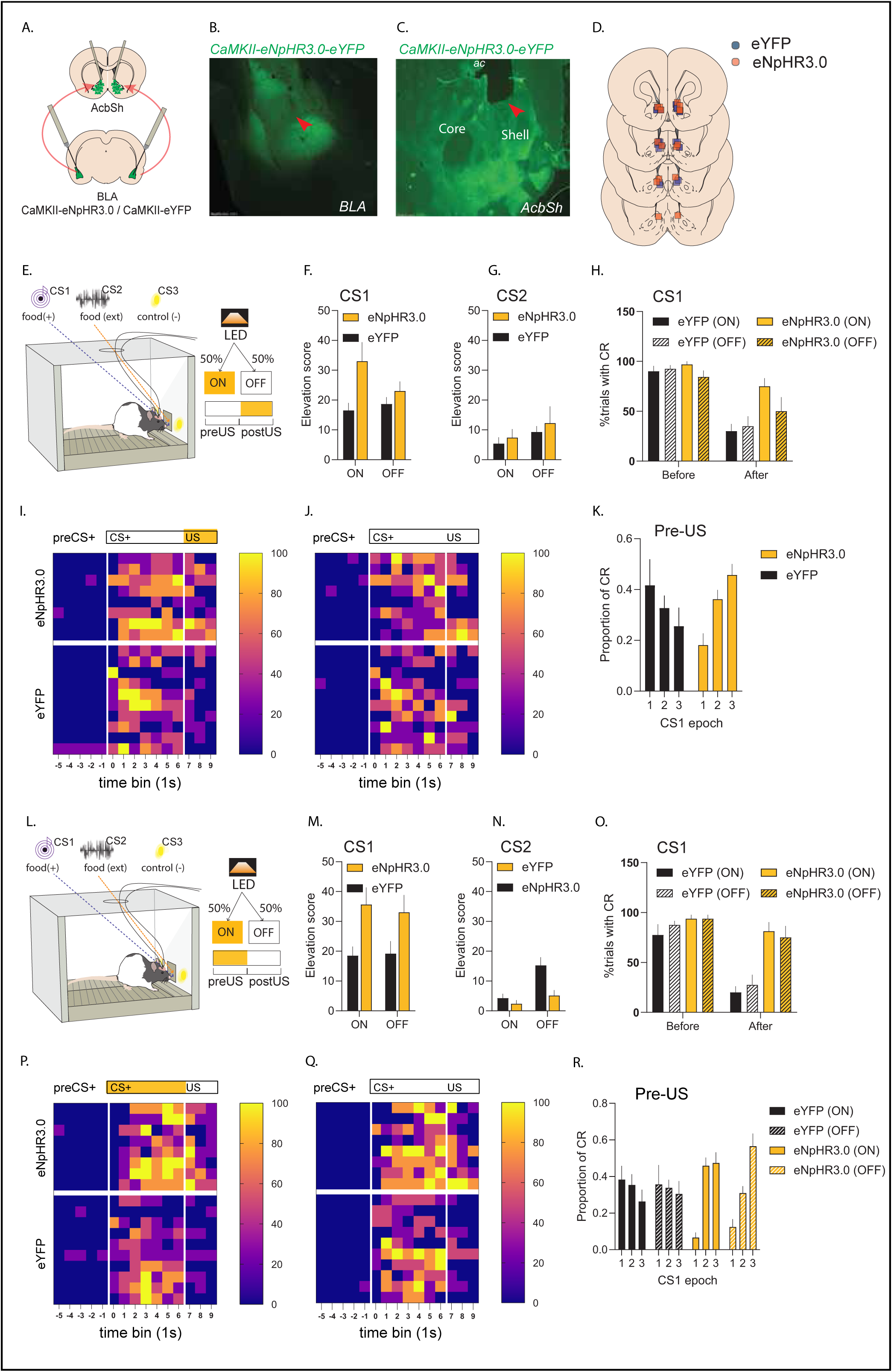
State-specific optogenetic silencing of BLA→AcbSh on appetitive transitions. ***A.*** Schematic of eYFP or eNpHR3.0 injected in BLA and fiber optic cannulae positioned above AcbSh. ***B.*** Representative photomicrograph of spread of eNpHR3.0-eYFP in BLA, *injection tip. ***C.*** Representative photomicrograph of BLA terminal labeling in accumbens, *cannula tip above AcbSh. ***D.*** Locations of fiber optic cannulae in AcbSh mapped on coronal sections. ***E.*** Schematic depicting appetitive test. CS1: food-reinforced; CS2: extinguished; CS3: never reinforced; LED delivered following US delivery (or equivalent onset for no-US CS2 and CS3) on 50% of trials of each CS. ***F-G.*** Mean CS-generated seeking responses for CS1 (***F.***) and CS2 (***G.***). ***H.*** Percentage of trials paired with seeking before and after US delivery for CS1. ***I-J.*** Heatplot depicting percentage of trials paired with seeking per 1s time bin (columns) per subject (rows) for CS1 with (***I.***) and without (***J.***) photoinhibition. ***K.*** Effects of post-US photoinhibition on shifts in seeking response following CS1 onset prior to US delivery. ***L.*** Schematic depicting appetitive test. CS1: food-reinforced; CS2: extinguished; CS3: never reinforced; LED delivered at CS onset and terminating at US delivery (or equivalent offset for no-US CS2 and CS3) on 50% of trials of each CS. ***M-N.*** Mean CS-generated seeking responses for CS1 (***M.***) and CS2 (***N.***). ***O.*** Percentage of trials paired with seeking before and after US delivery for CS1. ***P-Q.*** Heatplot depicting percentage of trials paired with seeking per 1s time bin (columns) per subject (rows) for CS1 with (***I.***) and without (***J.***) photoinhibition. ***R.*** Effects of pre-US photoinhibition on subsequent reward-mediated transitions in seeking.

Silencing BLA→AcbSh after reward delivery significantly increased overall seeking generated by the CS+ (CS1 group main effect: eNpHR3.0 vs eYFP, F_(1,16)_= 5.247, *p=*.036; **Figure 5F**). This difference was light-dependent (group x light, F_(1,16)_=5.352, *p*=.034; simple effects group difference for on and off trials were F_(1,16)_=.6.744, *p*=.019 and F_(1,16)_=1.252, p=.280 respectively; **Figure 5F**). It was also cue-dependent, since there were no significant group differences to the extinguished cue (CS2, group main effect: F_(1,16)_=.476, *p*=.5; group x light: F_(1,16)_=.028, *p*=.869, **Figure 5G**) or to the CS- (CS3, group main effect: F_(1,16)_=.626, *p*=.44; group x light: F_(1,16)_=.885, *p*=.361).

To determine whether BLA→AcbSh silencing impaired transitions driven by reward delivery, we confirmed that there was significant suppression of seeking following reward delivery. As before, the proportion of trials that elicited a seeking response was significantly higher before than after reward (main effect: before vs. after, F_1,16_=34.486, p=.000, **Figure 5H-J**). Critically, silencing BLA→AcbSh pathway only during the consummatory period (after reward) significantly promoted seeking within that period in a light-dependent manner (after reward, proportion of trials that elicited a seeking response: group x light, F_1,16_=5.818, p=.028; simple effects group On, F_1,16_=16.941, p=.001; group, Off, F_1,16_=.79, p=.387; before reward, group x light, F_1,16_=2.349, p=.145; simple effects group On, F_1,16_=1.019, p=.328; group, Off, F_1,16_=1.257, p=.279; **Figure 5H**). These findings show that BLA→AcbSh is required for the suppression of seeking following reward delivery.

We also inspected behavior prior to reward delivery to determine whether the effects of silencing of BLA→AcbSh during reward delivery were specific to transitions in the post-reward domain, or whether disruptions to this transition had consequences on appetitive behavior generally. While we did not find significant transitions prior to reward in this study, we found that the groups significantly differed in how they distributed seeking as the time of reward delivery approached. The distribution of active seeking was significantly linear in group eNpHR3.0 (F_(1,7)_=15.541, p=.006; **Figure 5K**) indicating that seeking peaked in the seconds prior to reward delivery. This trend was significantly different from group eYFP (group x linear trend, F_(1,16)_=6.161, p=.025; **Figure 5K**) and we confirmed that this difference was driven by group eNpHR3.0 allocating more seeking responses in the period most proximal to reward than group eYFP (group, epoch 3, F_(1,16)_=12.921, p=.002). Thus, silencing BLA→AcbSh pathway during reward delivery can significantly alter the topography of CS-generated seeking activity.

We tested the reverse condition, silencing the BLA→AcbSh pathway exclusively before reward delivery (**Figure 5L)**. This produced a significant increase in overall seeking during the CS+ (group main effect: eNpHR3.0 vs eYFP, F_(1,16)_= 6.961, *p=*.018; **Figure 5M**). However, this behavioral enhancement was not light-dependent (group x light, F_(1,16)=_ .294, p=.595). Nor was it specific to the CS+. Overall responding to the extinguished cue (CS2) was also enhanced (group main effect, F_(1,16)_=7.791, p=.013) although this enhancement was specific to trials without photoinhibition (group x light, F_(1,16)_=6.253, p=.023; simple effects, group(on), F_(1,16)_=.486, p=.496; simple effects, group(off), F_(1,16)_=15.463, p=.001), **Figure 5N**. We confirmed there was no baseline difference between groups (CS3, group main effect: F_(1,16)_= .024, *p=*.879; group x light, F_(1,16)_= 1.75, *p=*.204), nor were the groups different across CS1, CS2, or CS3 on the day prior to test (CS1 group main effect: F_(1,16)_= .702, *p=*.414; CS2 group main effect, F_(1,16)_=.017, *p=*.898; CS3 group main effect: F_(1,16)_= .181, *p=*.676).

Did BLA→AcbSh silencing before reward delivery affect the topography of CS-generated seeking prior to reward delivery? The topography of seeking prior to reward was significantly linear in group eNpHR3.0 (overall: F_(1,7)_=25.608, p=.000, **Figure 5R**) regardless of light condition (On condition: F_(1,7)_=26.518, p=0; Off condition: F_(1,7)_=16.619, p=.001; **Figure 5R**) indicating that overall, seeking peaked in the seconds prior to reward delivery. This trend was significantly different from group eYFP (group x linear trend, F_(1,16)_=8.255, p=.011) and we confirmed that this difference was driven by group eNpHR3.0 allocating more seeking responses in the period most proximal to reward (group, epoch 3, F_(1,16)_=8.086, p=.012) and fewer in the period most distal from reward (group, epoch 1, F_(1,16)_=7.165, p=.017) relative to group eYFP, **Figure 5R**. Thus, silencing BLA→AcbSh pathway before reward delivery significantly modified the topography of CS-generated seeking activity.

Finally, we inspected behavior after reward delivery. We confirmed that overall, seeking was significantly suppressed following reward delivery. That is, the proportion of trials that elicited a seeking response was significantly higher before reward delivery than after (main effect: before vs. after, F_1,16_=24.779, p=.000), **Figures 5O-Q**. Critically, silencing BLA→AcbSh pathway exclusively before reward delivery prevented this suppression (after reward, group main effect, F_1,16_=20.894, p=.000; simple effects group On, F_(1,16)_=32.584, p=.001; group, Off, F_(1,16)_=9.543, p=.007; before reward, group x light, F_1,16_=2.349, p=.145; simple effects group On, F_1,16_=1.019, p=.328; group, Off, F_1,16_=1.257, p=.279).

### BLA→AcbSh projections target LH-projecting AcbSh MSNs

To identify a putative output through which BLA→AcbSh might shape seeking behavior, we focused on the AcbSh projections to the LH, given their role in gating consummatory and approach behaviors (Gibson et al., 2018; O’Connor et al., 2015; Stratford & Kelley, 1999). We asked whether BLA neurons synapse onto LH-projecting AcbSh neurons. BLA neurons were transduced with ChR2 and LH-projecting AcbSh neurons were identified by injection of the red-fluorescent retrograde tracer CTb-555 into the LH. Acute slices containing ChR2-expressing terminals from BA and AcbSh→LH labelled neurons were prepared for whole-cell patch-clamp recordings. Brief light stimulation (3-5ms) of BLA terminals evoked an excitatory post synaptic current (EPSC) in 7 of the 13 neurons recorded (**Figures 6A-C**). The onset of EPSC was 5.3 ± 0.6 ms after the light onset, indicative of a monosynaptic response.

**Figure 6.**
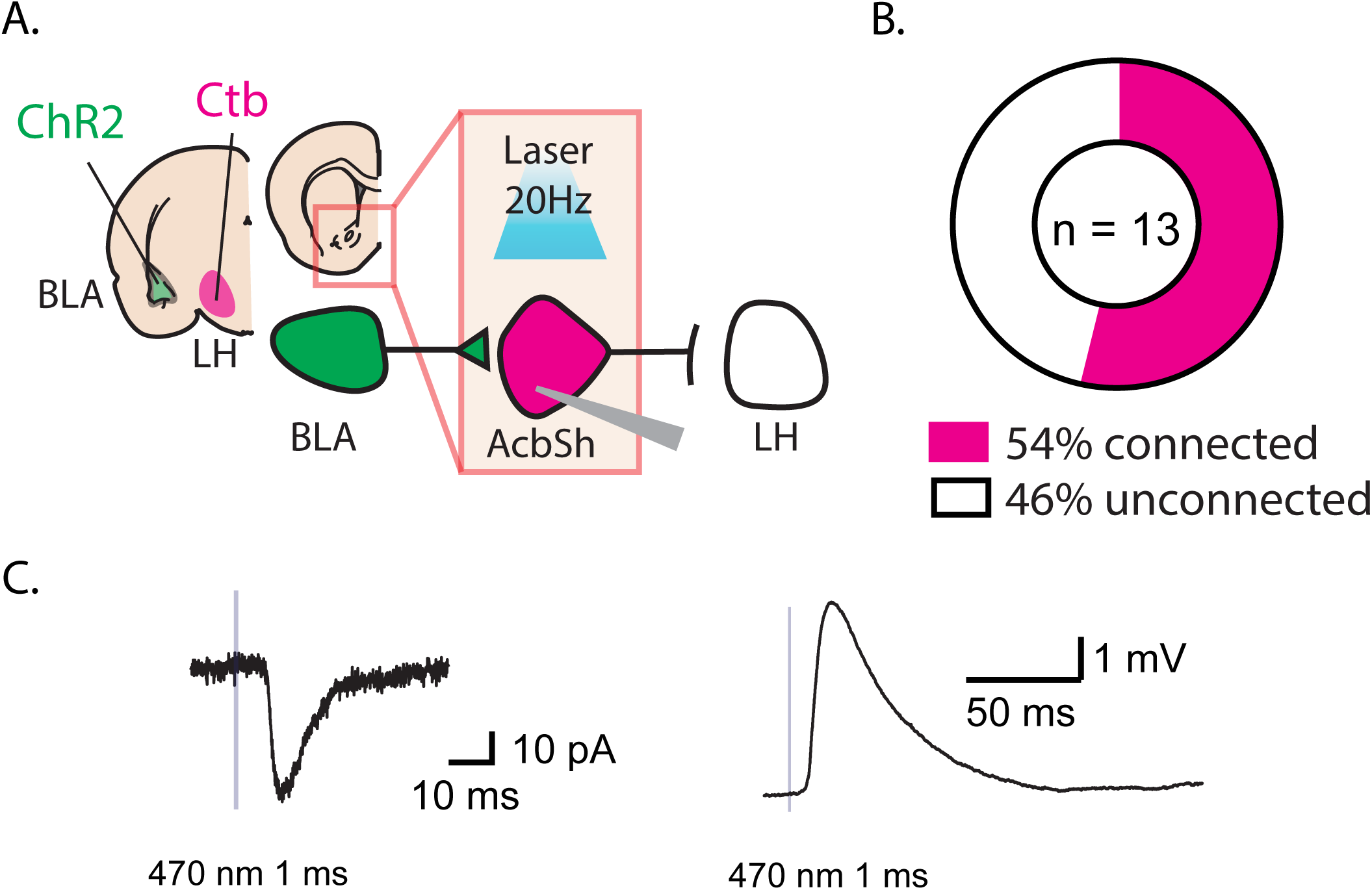
Characterization of LH-projecting AcbSh neurons. ***A.*** Schematic of combined tract tracing with optogenetic stimulation to identify LH-projecting AcbSh neurons responsive to photostimulation of BLA terminals. ***B.*** Percentage of recorded LH-projecting AcbSh cells responding with an EPSC following optogenetic-mediated stimulation of BLA-terminals. ***C.*** Representative EPSC (left panel) and EPSP (right panel) in a connected cell following a brief light (1 ms) stimulation of BLA terminals. *N*=13

## General Discussion

Behavioral transitions are critical for flexible and responsive sequences of behavior. They require not only the initiation, but also the termination of actions. For example, a variety of behavioral forms (e.g., approach, nose around, grab) are initiated to seek goals such as food; and other forms quickly replace these at goal receipt (e.g., ingesting, handling, hoarding) (Timberlake, 1994). Here, we sought to understand the causes and mechanisms for these appetitive transitions. We show that reward delivery is a critical trigger for the switch from seeking to taking. Presentations of the appetitively-conditioned CS (CS+) robustly induced seeking that was in turn abruptly and transiently reduced during reward delivery. This reduction was not due to response fatigue because seeking persisted during an equivalent period when reward was omitted under full or partial reward omission schedules. Nor was it due to satiety as it was immediately followed by a period of recovered seeking. Critically, we show that the functional operation required for these transitions is active inhibition of seeking mediated via a BLA→AcbSh pathway. Also, that the processes targeted by this inhibition are linked to and have consequential effects on the appetitive state. Finally, that LH-projecting AcbSh cells receive monosynaptic BLA input, indicating that the BLA→AcbSh pathway is well-positioned to regulate appetitive seeking behavior. Together, these findings implicate a limbic-striato-hypothalamic circuit in the sculpting of appetitive sequences.

### BLA→AcbSh pathway sculpts rapid downward transitions in appetitive behavior

The sculpting of action sequences is largely considered a domain of cortico-basal ganglia circuits (Jin & Costa, 2015), particularly involving dorsal striatopallidal pathways in stopping previous actions in order to promote new actions (Tecuapetla, Jin, Lima, & Costa, 2016). However, involvement of ventral striatal circuitry as a “task sequencing device” have also been proposed based on early observations in cocaine-seeking rats that nucleus accumbens neurons are often responsive at the termination of action segments (Chang, Sawyer, Lee, & Woodward, 1994; Woodward, Janak, & Chang, 1998). Our findings support this early proposition for ventral striatum and suggest that a BLA→AcbSh pathway is required for appetitive seeking-taking sequences. We show that this role involves the rapid inhibition of previous seeking actions as animals transition to reward taking. These findings are consistent with more recent work implicating BLA-AcbSh interactions in constraining appetitive behavior under a variety of conditions: by cues that interrupt ongoing appetitive behavior, by an unexpected distracting stimulus, stimuli that signal no reward, or punishment; and by conditions of extinction (non-reinforcement) and satiety that occur over longer timescales (Lafferty et al., 2020; Millan & McNally, 2011; O’Connor et al., 2015; Piantadosi, Yeates, Wilkins, & Floresco, 2017; Tye, Cone, Schairer, & Janak, 2010). However, our findings also demonstrate several critical features of this regulation.

First, BLA→AcbSh pathway contributions promote a specific behavioral form – they are inhibitory – and they require an active appetitive state. Indeed, there was no effect of silencing on responding to an extinguished cue or to a cue never paired with reward. Thus, silencing this pathway did not simply potentiate responding or generate an appetitive state. It also had no effect on performance against a high background of seeking generated at the onset of the appetitive CS, prior to reward. Instead, two points of inflection in behavior were sensitive to disruption following BLA→AcbSh pathway silencing. The first was at the time of reward delivery. The second was in the seconds prior to reward delivery, following the peak in CS-generated seeking. This suggests that dynamic fluctuations in CS-generated seeking are governed by the BLA→AcbSh pathway.

Second, the regulation of seeking by a BLA→AcbSh pathway was process-specific because it was not required to suppress responding to an extinguished CS. This finding is important given previous findings that extinction of appetitive behavior depends on AcbSh (Millan et al., 2010) and its interactions with BLA (Millan & McNally, 2011). However, these findings depended on instrumental rather than Pavlovian responding, and the suppression of seeking by extinction may involve separate systems depending on the mode of original learning (Bouton, Maren, & McNally, in press). Certainly active inhibition of appetitive responding is not itself sufficient for engagement of the BLA→AcbSh pathway.

What then are the conditions for BLA→AcbSh recruitment? Previous studies have suggested that reward unavailability may be essential (Lafferty et al., 2020). However, our findings suggest that reward unavailability, whether signaled by an extinguished cue or a never rewarded cue, was not itself sufficient to reveal a role for the BLA→AcbSh in sculpting behavior. Yet, AcbSh and its pathways control appetitive seeking under rather diverse conditions, including by stimuli interrupting a maintained instrumental or unconditioned appetitive response, stimuli that signal punishment, and during stimulus-driven choice (Lafferty et al., 2020; Leung & Balleine, 2013; O’Connor et al., 2015; Piantadosi et al., 2017). Together with these findings, we suggest that the diverse conditions under which AcbSh pathways regulate performance may necessarily involve conditions requiring rapid or dynamic shifts in behaviors surrounding appetitive transitions.

Third, our findings demonstrate the regulation of seeking was anatomically specific to the BLA→AcbSh pathway; silencing BLA→AcbC was largely ineffective and had only a modest effect, increasing the vigor (rate) of seeking during reward delivery. This is consistent with previous reports indicating that pharmacological inactivation of AcbC can have disinhibitory effects on unproductive appetitive responding, and that these effects are modest relative to similar manipulations targeting the AcbSh (Ambroggi et al., 2011). Moreover, the observed functional distinctions between BLA→AcbSh and BLA→AcbC are in line with anatomically segregated projection profiles (Heimer et al., 1997; Heimer, Zahm, Churchill, Kalivas, & Wohltmann, 1991; Zahm & Heimer, 1993) and afferents from BLA, where inputs are largely organized along the anterior-posterior axis of the BLA (Hamlin, Clemens, Choi, & McNally, 2009; Heimer et al., 1997; Wright et al., 1996). Here, we used a dual-labeling retrograde tract tracing approach targeting dorsal AcbSh and AcbC-projecting BLA pathways. We found these pathways to be highly segregated within BLA with few overlapping cells, and topographically defined along two axes: the anterior-posterior plane of the BLA, and between basal (BA) and lateral (LA) nuclei of the amygdala. That is, AcbSh was preferentially innervated by posterior BA while AcbC was preferentially innervated by anterior BLA, across both BA and LA. These findings suggest that posterior BLA may be particularly implicated in the rapid regulation of seeking. Indeed, previous findings suggest that posterior and anterior BLA are preferentially recruited by rewarding and aversive stimuli respectively (Kim, Pignatelli, Xu, Itohara, & Tonegawa, 2016); have genetically-distinct pyramidal neurons (Kim et al., 2016); are functional distinct (Kantak, Black, Valencia, Green-Jordan, & Eichenbaum, 2002; McLaughlin & Floresco, 2007; Millan et al., 2015); and critically, that posterior BLA can exert regulatory control over some forms of appetitive behavior (McLaughlin & Floresco, 2007; Millan et al., 2015).

Finally, using state-dependent silencing of the BLA→AcbSh pathway, (i.e. after or before reward delivery), our findings demonstrate that BLA→AcbSh control over seeking targets specific processes that have consequences for ongoing appetitive sequences. We found that silencing after but not before reward delivery impaired the dynamic changes in appetitive seeking both prior to and after delivery. In the reverse condition, silencing before but not after reward delivery, produced a similar trend, though the effects were more generalized extending even to trials that were not paired with photoinhibition. Regardless, the pattern of effects suggests a co-dependency between CS- and US-generated behavior and that the properties required for this co-dependency are targeted by BLA→AcbSh. What these properties are remain to be defined, although they may be related to sensory features of the US (Shiflett & Balleine, 2010) and given that for appetitively conditioned cues, proximity to ingestion is increasingly controlled by sensory properties of the US (Galarce, Crombag, & Holland, 2007). Moreover, co-dependencies between CS- and US-generated behavior are assumed in associative learning models (A. Wagner, 1981; A. R. Wagner & Donegan, 1989) and their neurobiological instantiation will be of interest in future studies.

### AcbSh as the interface to behavioral control columns

Learned appetitive behaviors depend on close functional links with preorganised modules for motivational and motor control (Timberlake, 1994) and their neural instantiation in behavioral control columns involving the hypothalamus and brainstem (Swanson, 2000). Indeed, LH has distinct neuronal populations encoding appetitive versus consummatory behavior (Jennings et al., 2013), and subpopulations of LH neurons can control feeding in different ways (Marino et al., 2020). The AcbSh provides the major source of striatal input to the LH. Notably, these inputs arise largely from D1- than D2-expressings MSNs, targeting GABAergic ensembles within LH (Gibson et al., 2018; O’Connor et al., 2015). Here we determined that more than half of detected LH-projecting AcbSh cells receive functional contacts from BLA, confirming a BLA→AcbSh→LH circuit. They suggest that BLA mediated stimulation of LH-projecting D1 MSNs in AcbSh may be a viable route through which seeking is rapidly terminated at the transition from seeking to taking.

LH, particularly its anterior division, is anatomically well positioned to modulate seeking activity given its access across critical nodes of motivated behavior. These include key substrates of the behavioral state control system, via LH outputs to midbrain regions–ventral tegmental area (VTA), its medial inter-fascicular nucleus (IF), as well as dorsal raphe (DR); metabolism and feeding via outputs to bed nucleus stria terminalis anteromedial group (BSTamg), anterodorsal preoptic nucleus, and peri-locus coeruleus; and fight or flight defensive behavior via outputs to dorsal premammilary nucleus (PMd) and periaqueductal grey area (PAG) (Marino et al., 2020; Nieh et al., 2015; Thompson & Swanson, 2010). In addition to these three descending pathways, outputs of the LH also provide an ascending relay back into top-down control over motivated behavior via cortical-striatal-thalamic-hypothalamic loops involving LH projections to midline paraventricular thalamus (PVT) and infralimbic prefrontal cortex (ILPFC), and from here, a return to AcbSh (Bouton et al., in press; Thompson & Swanson, 2010). Among these relays, LH GABAergic projections targeting VTA promote approach to food and eating as well as sufficient reinforcement to sustain place preference, intracranial self-stimulation, social interactions, and exploratory behaviors (Marino et al., 2020; Nieh et al., 2015). LH GABAergic neurons also control both approach and eating via projections to a relatively undefined brainstem area medial of LC (Marino et al., 2020; Nieh et al., 2015). It will be of interest whether BLA inputs to AcbSh recruits selectively among these LH routes or has a broader network-like impact on the outputs of the LH (Millan et al., 2010; Stratford & Kelley, 1999).

### Concluding Remarks

We show that appetitive behavioral transitions are actively controlled at the limbic-striatal interface, substantiating early indications of the accumbens in sequencing behaviors around appetitive rewards (Chang et al., 1994; Woodward et al., 1998). Reward receipt is a proximal trigger for transition between reward seeking and taking and active inhibition is the functional operation for this transition. The BLA→AcbSh pathway is critical to this functional operation, likely via its interface with behavioral control columns in the LH. We suggest that the BLA→AcbSh pathway is a core component of an appetitive switching system, recruited under conditions requiring rapid or dynamic shifts in appetitive behavior, and that this pathway enables these shifts by actively inhibiting seeking. It remains to be determined whether this system is required for seeking-taking transitions of non-ingested rewards such as cocaine, where seeking responses are similarly terminated by the receipt of the reward (Chang et al., 1994) and given the role of the AcbSh in the regulation of cocaine-seeking behaviors (Peters et al., 2008; Sutton et al., 2003). A possibility is that compromises to this system may contribute to the heightened and persistent seeking characteristic of conditions such as addiction.

## Acknowledgments

Data reported here are archived in the UNSW Long Term Data Archive (ID:). This work was supported by the Australian Research Council (DP200102576) and National Health and Medical Research Council (GNT1138062).

